# Benchmarking of deep neural networks for predicting personal gene expression from DNA sequence highlights shortcomings

**DOI:** 10.1101/2023.03.16.532969

**Authors:** Alexander Sasse, Bernard Ng, Anna E. Spiro, Shinya Tasaki, David A. Bennett, Christopher Gaiteri, Philip L. De Jager, Maria Chikina, Sara Mostafavi

## Abstract

**Introductory Paragraph**

Deep learning methods have recently become the state-of-the-art in a variety of regulatory genomic tasks^1–6^ including the prediction of gene expression from genomic DNA. As such, these methods promise to serve as important tools in interpreting the full spectrum of genetic variation observed in personal genomes. Previous evaluation strategies have assessed their predictions of gene expression across genomic regions, however, systematic benchmarking is lacking to assess their predictions across individuals, which would directly evaluates their utility as personal DNA interpreters. We used paired Whole Genome Sequencing and gene expression from 839 individuals in the ROSMAP study^7^ to evaluate the ability of current methods to predict gene expression variation across individuals at varied loci. Our approach identifies a limitation of current methods to correctly predict the direction of variant effects. We show that this limitation stems from insufficiently learnt sequence motif grammar, and suggest new model training strategies to improve performance.

## Main

Sequence-based deep learning methods are emerging as powerful tools for a variety of functional genomic prediction tasks. These models take as input genomic DNA, and learn to predict context-dependent functional outputs like transcription factor binding^2,8,9^, chromatin state^10–13^ and gene expression values^1,14^. State of the art models can reproduce experimental measurements with a high degree of accuracy and enable mechanistic insights through their learnt DNA features^1,2,15^. Yet, the true potential of these sequence-based models lies in their ability to predict outcomes for arbitrary sequence inputs – a space too large for experimental methods to fully explore. While partial evaluations through expression quantitative trait loci (eQTL)^1,16^ studies or massively parallel reporter assays (MPRA)^17^ have shown promise, the broader application of these models as personalized DNA interpreters has not been comprehensively assessed. We address this by conducting an extensive analysis using paired Whole Genome Sequencing (WGS) and cerebral-cortex RNA-sequencing data from the ROSMAP datasets^7^ with measurements from 839 individuals. Our study bridges the gap between the known potential and the actual performance of these models in personalized genomics interpretation.

To start, we focus our evaluation on Enformer^1^, the top-performing deep learning model. Enformer is trained to predict various functional outputs from (*cis*) sub-sequences from the Reference genome. This training approach allows Enformer and other deep learning models to identify short DNA sub-sequences (motifs) that are shared across the genome and exploits variations in motif combinations across genomic regions to make context-dependent predictions. As a control experiment, we used the pre-trained Enfomer model, provided it with sub-sequences around the TSS from the Reference genome and evaluated its predictions on population-average gene expression (n=13,397 expressed protein coding genes) from the cerebral cortex (**Fig. 1A-B**). To account for the differences between data types that were used during Enformer’s training and our study, we used a fine-tunning strategy, whereby we trained an elastic net model on top of the predictions from Enformer’s output tracks (see Methods). Consistent with the expectation for this type of evaluation, we observed good prediction accuracy as measured by the Pearson correlation coefficient R=0.58 (**Fig. 1B**). The results were similar when we restricted the analysis to a smaller set of genes (n=3,401) overlapping Enformer’s test regions (R=0.51; **Supplementary Fig. 1**).

**Figure 1.**
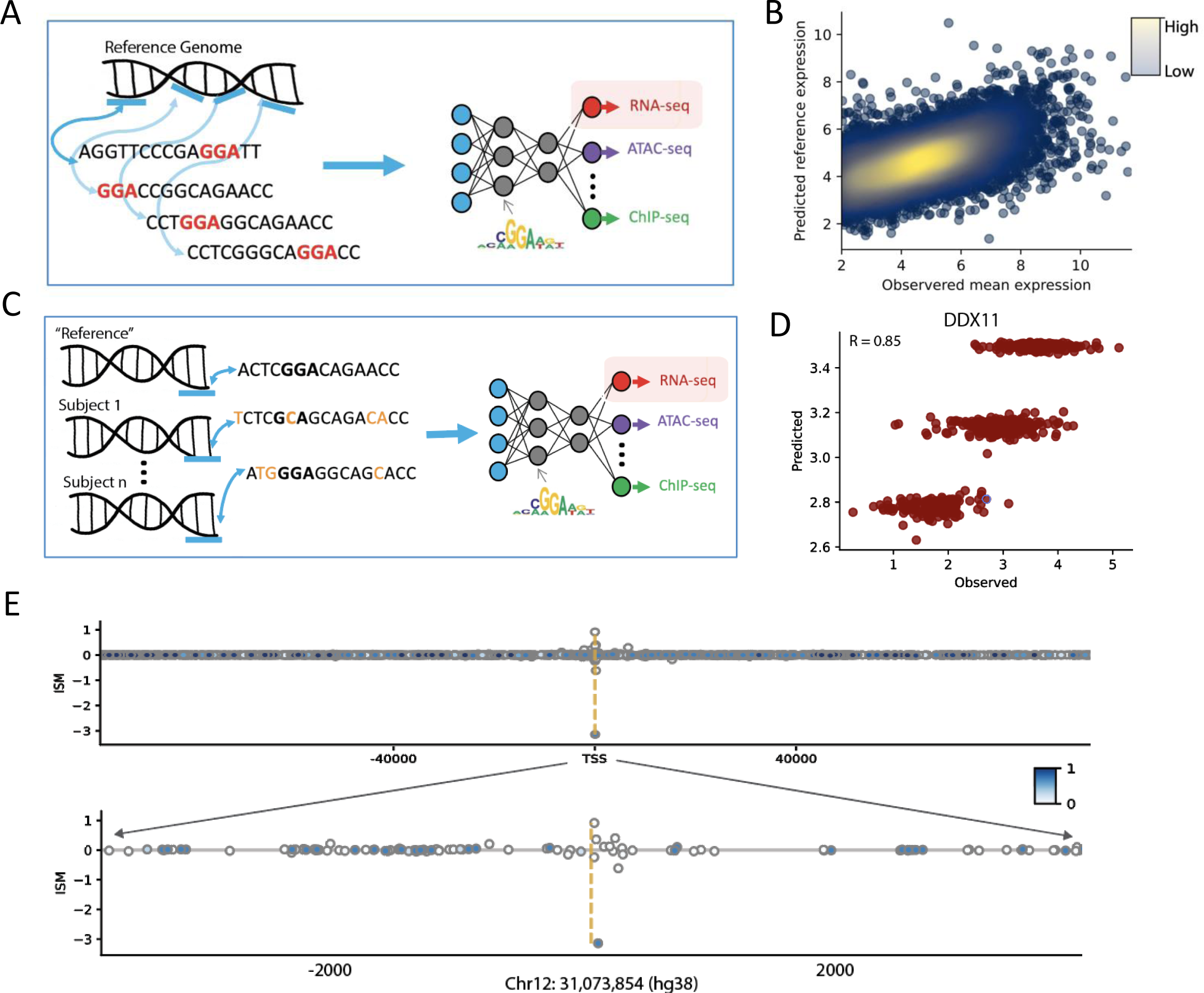
Evaluation of Enformer across genomic regions and select loci. (A) Schematic of the Reference-based training approach. Different genomic regions from the Reference genome are treated as data points. Genomic DNA underlying a given region is the input to the model, and the model learns to predict various functional properties including gene expression (CAGE-seq), chromatin accessibility (ATAC-Seq), or TF binding (ChIP-Seq). (B) Population-average gene expression levels in cerebral cortex (averaged in ROSMAP samples, n=839) for expressed genes (n=13,397) versus Enformer’s predictions. (C) Schematic of the per-locus evaluation strategy. (D) Predicted and observed DDX11 gene expression levels in cortex for individuals in the ROSMAP cohort (n=839). Each dot represents an individual. Output of Enformer is fine-tuned using an elastic net model (Methods). (E) In-silico mutagenesis (ISM) values for all SNVs which occur at least once in 839 genomes within 98Kb of *DDX11* TSS. SNVs are colored by minor allele frequency (MAF).

While Enformer is not explicitly trained on genetic variation data, once trained, it holds promise that it has learnt the *cis* regulatory logic of gene expression and so can predict the impact of arbitrary genetic variation on its outputs. To evaluate its performance in this setting, which is distinct from the cross-genome performance evaluated above, we applied Enformer to predict individual-specific gene expression levels based on personal genomic sequences (Methods; **Fig. 1C**). As a positive example, we first present here results for a highly heritable gene (heritability *r*^2^=0.8) *DDX11*. *DDX11*’s variance in expression across individuals can be attributed to a single causal single-nucleotide variant (SNV) using statistical fine-mapping^16^. Using WGS data, we created 839 input sequences of length 196,608bp centered at the transcription start site (TSS), one per individual for the gene (**Fig. 1C, Supplementary Fig. 2**, Methods). Applying Enformer to these input sequences we observed a Pearson correlation of 0.85 (p<1e-200) between predicted and observed gene expression levels across individuals (**Fig. 1D**). Further, *in-silico* mutagenesis (ISM) at this locus showed that Enformer utilized a single SNV with high correlation to gene expression (eQTLs) in making its predictions (**Fig. 1E**). This SNV is the same causal SNV that was identified through statistical fine-mapping with Susie^16^. Thus, at this locus, Enformer is able to identify the causal SNV amongst all those in LD, and in addition provides hypotheses about the underlying functional cause, in this case the extension of a repressive motif (**Supplementary Fig. 3**).

However, the impressive predictions on *DDX11* proved to be the exception rather than the rule. When we tested 6,825 cortex-expressed genes, we found a large distribution in Pearson’s *R* (**Fig. 2A, Supplementary Table 1**; min R=-0.76, max R=0.84, mean = 0.01). Surprisingly, while the predictions were significantly correlated to observed expression for 598 genes (*FDR_BH_=0.05*, Methods), they were significantly anti-correlated with the true gene expression for 195 (33%) of these genes. For example, predicted *GSTM3* gene expression values are anti-correlated with the observed values (R= -0.49; p<1e-200, **Fig. 2B**). We performed several sensitivity analyses to which these results proved robust (Methods, **Extended Data Fig. 1**): these results are not sensitive to output track fine-tuning, or model ensembling as done in Enformer, or subseting the analysis to a smaller set of genes that have easily detectable causal variants based on statistical fine-mapping (**Supplementary Table 1**). Overall, these results imply that the model fails to correctly attribute the variants’ direction of effect (*i.e.,* whether a given variant decreases or increases gene expression level).

**Figure 2.**
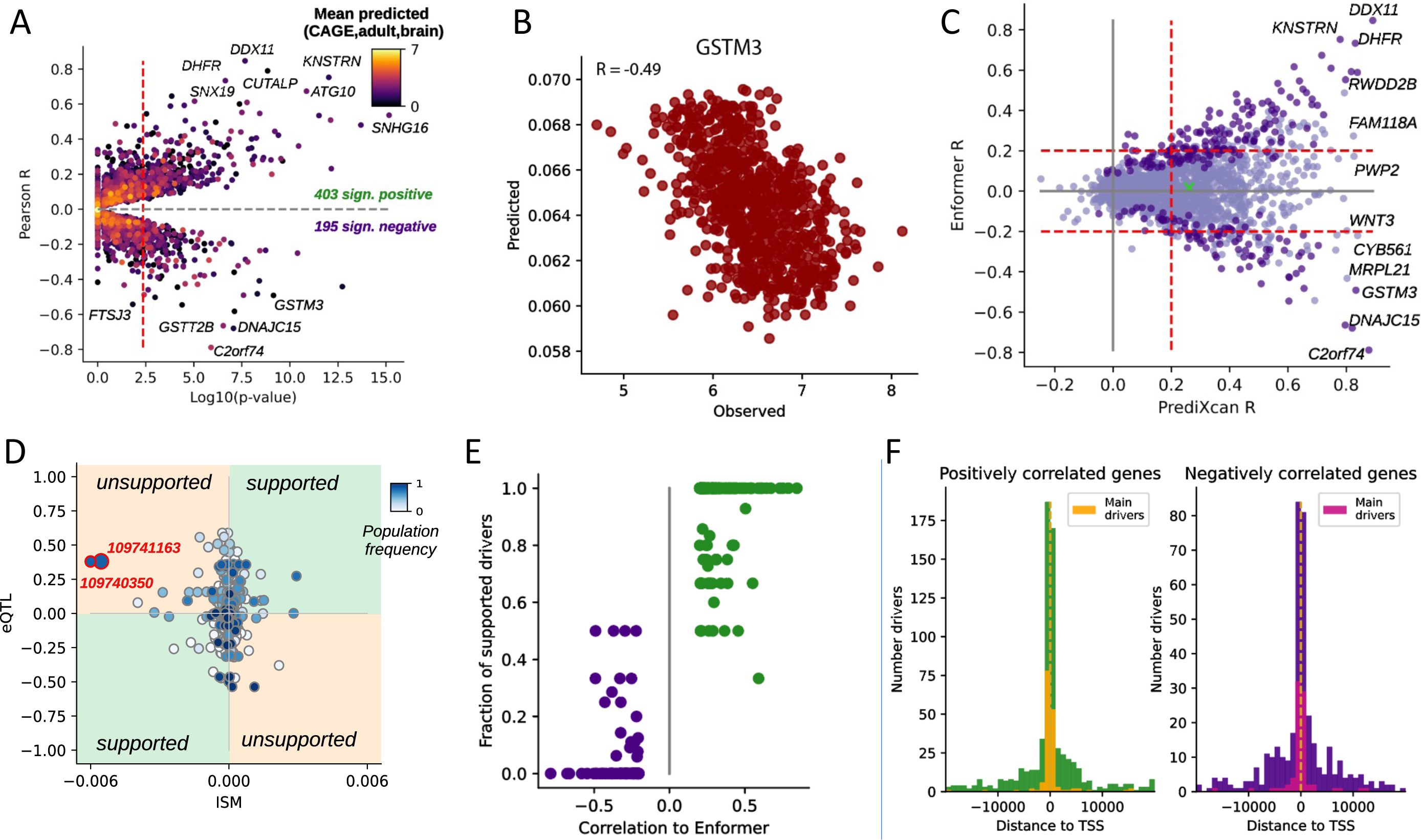
Evaluation of Enformer on prediction of gene expression across individuals. (A) Y-axis shows the Pearson R coefficient between observed expression values and Enformer’s predicted values per-gene (genes=6,825, individuals=839). X-axis shows the negative log10 p-value, computed using a gene-specific null model (Method, one-sided T-test, permutation analysis with n=50 independent samples per gene). The color represents the predicted mean expression using the most relevant Enformer output track (“CAGE, adult, brain”). Red dashed line indicates FDR_BH_=0.05. (B) Y-axis shows the prediction from Enformer’s “CAGE, adult, brain” track across individuals for the *GSTM3* gene (n=839), x-axis shows the observed gene expression values. (C) Pearson R coefficients between PrediXcan predicted versus observed expression across individuals is shown on the x-axis, Enformer’s Pearson R coefficients are shown on the y-axis. Red lines indicate threshold for significance (abs(R)>0.2, Bonferroni corrected nominal p-value), darker colored dots are significant genes from panel A. Green cross represents the location of the mean across all x- and y-values. (D) ISM value versus eQTL effect size for all SNVs (n=706 with MAF>0.1) within the 196Kb input sequence of the *GSTM3* gene. Red circles represent driver SNVs. SNVs are defined as supported or unsupported based on the concordance with the sign of the eQTL effect size. (E) Fraction of supported driver SNVs per gene (y-axis) versus Pearson’s R coefficients between Enformer’s predictions and observed expressions (x-axis) (n=87 supported genes, n=161 unsupported genes). (F) Number of driver SNVs within the 1000bp window of the TSS. Main drivers are the drivers with the strongest impact on linear approximation, shown in different colors. Left plot, n=983 driver SNVs; Right plot, n=564 driver SNVs.

We then compared Enformer against a widely-used linear approach called PrediXcan^18^. PrediXcan constructs an elastic net model per gene from *cis* genotype SNVs across individuals. Unlike Enformer, PrediXcan is explicitly trained to predict gene expression from variants but it does not take into account variants that were not present in its training data and cannot output a prediction for unseen variants. While the models are conceptually different, the PrediXcan model gives a lower bound on the fraction of gene expression variance that can be predicted from genotype. Further, genes that are significantly predicted with PrediXcan should have at least one causal variant somewhere in the genomic region used for the predictions and thus provide a substantial set of loci for assessing Enformer’s predictions. We used the previously published prediXcan model that was trained on GTEx cortex^18^ and simply applied it to ROSMAP samples. Hence neither Enformer nor PrediXcan have seen the ROSMAP samples prior to their application. As shown in **Fig. 2C**, for the 1,570 genes where PrediXcan’s elastic net model was available, performance of Enformer is substantially lower (921 significantly predicted gene by PrediXcan vs. 162 by Enformer, Mean R Enformer = 0.02, Mean R PrediXcan = 0.26**, Supplementary Table 1**). Further, PrediXcan did not have the same challenge with mis-prediction of the direction of SNV effect (*i.e.,* all PrediXcan’s significantly predicted genes have a positive correlation between predicted and observed). When we ignore the sign of the Enformer’s correlation values, we observe that both models, despite their conceptual differences, show some predictive power for the same genes (R=0.58, **Supplementary Fig. 4**). This supports the observation that Enformer can identify genes whereby genetic variation across individuals significantly impacts gene expression values, but unlike PrediXcan, it is not able to determine the sign of SNV effects accurately. We note that Enformer predictions were evaluated against eQTLs in the original study using signed linkage disequilibrium profile (SLDP) regression^1,19^ demonstrating improved performance over competing models in terms of z-score, however, this previous result is based on evaluation across the genome, and not loci specific as we report here.

To investigate if these observations are specific to Enformer or more broadly apply to sequence-based deep learning models that follow the same training recipe, we trained a simple CNN that takes as input sub-sequences from the Reference genome centered at gene TSSs (40Kbp) and predicts population-average RNA-seq gene expression from cortex as output (Methods). This CNN can predict population-average gene expression in cortex with a similar accuracy as Enformer (R=0.57, **Extended Data Fig. 2A**), yet it has the same challenge with direction of the predictions across individuals (**Extended Data Fig. 2B**). Thus, our results on Enformer are likely to generalize to other sequence-based deep learning models trained in the same way.

To explore the causes for the negative correlation between Enformer predictions and the observed gene expression values we used two explainable AI approaches: ISM and input-Gradient (Supplementary Methods S2). These approaches approximate the output of a nonlinear neural network with a linear function that weights the contribution of each SNV through a process referred to as feature attribution. First, we confirmed that this approximation was reasonable for 95% of the examined genes (**Supplementary Fig. 5 and 6**). For each gene, based on its ISM attributions, we determined the main SNV driver(s) that dominate the differential gene expression predictions across individuals (Supplementary Methods S2). Across the 256 examined genes, we found that 32% have a single SNV driver, and the vast majority (85%) have five or fewer drivers (**Supplementary Fig. 7, Supplementary Table 2**) which determine the direction and correlation with the observed expression values. To understand how these driver SNVs cause mispredictions, we classified Enformer-identified driver SNVs into “supported” and “unsupported” categories based on the agreement of SNVs’ ISM attribution sign with the direction of effect according to the eQTL analysis (Methods). For this analysis, we computed marginal eQTL effect sizes, which do not distinguish causal variants from others in LD. However, it is important to note that the Enformer model is entirely agnostic to LD structure as it was trained with a single Reference genome. As such, Enformer predictions by construction assume a causal interpretation of the identified drivers variants. Thus, a comparison of Enformer-identified driver variants is informative because a sign discordance between the two strongly suggests that the Enformer effect is incorrect. On the other hand, the reverse analysis is not interpretable: an eQTL with a large marginal effect can have a low Enformer effect because it is not causal. As an example of sign discordance analysis, *GSTM3* has two common driver SNVs identified by Enformer yet their predicted direction of effect was unsupported based on the SNVs signed eQTL effect size (**Fig. 2D**). For all 256 inspected genes, we found that mispredicted genes had almost exclusively unsupported driver SNVs whereas correctly predicted genes indeed had supported driver SNVs **(Fig. 2E**). This analysis thus confirms that this small number of driver SNVs per gene are the cause of Enformer’s misprediction for the sign of the effect.

To investigate whether these unsupported attributions are caused by systematically erroneous sequence-based motifs that Enformer learns from the training data, we analyzed the genomic sequences around driver SNVs. We did not find any enrichment for specific sequence motifs (**Supplementary Fig. 8**). When we plotted the location of SNV drivers along the input sequences, we found that most drivers were located close to the TSS (**Fig. 2F, Supplementary Fig. 9 and 10**, Supplementary Methods S3), supporting a recent report^17^ that shows current sequence-based deep learning models mainly predict gene expression from genomic DNA close to TSS, despite using larger input DNA sequences. Further, when we analyzed ISM values in windows around the driver SNVs, we observed that the majority do not fall into coherent *attributional motifs* (short regions of sequence with consistent attribution) as would be expected if the model was picking up on biologically meaningful regulatory mechanisms (**Supplementary Fig. 11, Supplementary Table 3,** Supplementary Methods S4).

In summary, our results suggest that current sequence-based deep learning models trained on the input-output pair of a single Reference genome often fail to correctly predict the direction of SNV effects on gene expression. We further show that current neural network models perform worse than simple baseline approaches like PrediXcan in predicting the impact of genetic variation across individuals. For future development, we recommend that new models are not only assessed on genome-wide statistics of absolute causal eQTL effect sizes but also on a per-gene agreement between the sign and the size of the predicted and measured effect of causal variants.

We hypothesize that two complementary strategies will be fruitful for improving the prediction of gene expression across individuals. Firstly, current models are trained on sequences from a single Reference genome and learn sequence features that explain gene-to-gene expression variation, and thus have not been explicitly trained to learn how loci-dependent genetic variation impacts gene expression. The mechanisms that explain gene-to-gene variation may be distinct from those that explain interpersonal variation, for example, while promoter logic is important to determine which genes are expressed within a cell type, long-range interaction appears to be much more important for interpersonal variation^17^. Thus, training on input-outputs-pairs of diverse genomes and their corresponding gene expression measurements may be a way to increase sequence variation and learn these effects for accurate personalized predictions. Second, current methods do not accurately model all of the biochemical processes that determine RNA abundance. For example, post-transcription RNA processing (whose dependence on sequence is mediated via RNA-protein or RNA-RNA interactions) is entirely ignored. While including data sets that explicitly measure post-transcriptional regulatory processes and long-range interaction may improve modeling of these effects^4,6^, it is also possible that with sufficiently large paired WGS and gene expression training datasets, the resulting models will implicitly learn these mechanisms as long as they impact gene expression variation across individuals.

## Acknowledgements

We thank David R. Kelley for helpful comments on this manuscript. We thank the participants of ROS and MAP for their essential contributions and gift to this project. This work has been supported by many different NIH grants: P30AG10161 (to DAB), P30AG72975 (to DAB), R01AG15819 (to DAB), R01AG17917 (to DAB), U01AG46152 (to DAB and PLD), U01AG61356 (to DAB and PLD), R01AG057911 (to CG), R01AG06179 (to CG), R01AG036836 (to PLD), as well as a CIFAR research fellowship and an NSERC Discovery Grants (to SM). The funders had no role in study design, data collection and analysis, decision to publish or preparation of the manuscript.

## Author Contributions Statement

Conceived the study: SM, MC. Study design: SM, AS, MC. Data generation and quality control analyses: BN, AES, CG, PLD, ST, DAB. Analyses and interpretation: AS, AES, BN, SM, MC. Wrote the initial draft: SM, AS, BN. Read and provided comments on the manuscript: MC, BN, AES, PLD, CG, ST, DAB. Supervised the project: SM, MC.

## Methods

No specific ethics approval was needed to conduct the current study.

### WGS and RNA-seq datasets

We used n=839 subjects with available WGS (blood) and RNA-seq (cerebral cortex) from the ROS and MAP cohort studies^20^ (previously described^21^, also see Supplementary Methods). The 839 samples are from distinct individuals. Both studies were approved by an Institutional Review Board of Rush University Medical Center. All participants signed an informed and repository consent and an Anatomic Gift Act. Besides the availability of both WGS and cortex RNA-seq after pre-processing, no other exclusion criteria were used.

### Predicting gene expression with Enformer

#### Population-average gene expression

We centered the Reference genome (GRCh38) around the gene’s TSS (Gencode v27), and extracted the genomic sequence in the +/- 196,608 bp window, which was then used as input to Enformer (April 2022, v1). We performed this analysis for 13,397 brain expressed genes (for computational reasons, a random set of 6,825 genes among these were used in per-individual analyses described below). To use the outputs of Enformer predictions, we closely followed previous methodology. Specifically, for a given input sequence, Enformer makes predictions for 5,313 human output tracks and 986 bins. The predictions were obtained for all the 5,313 human output tracks, as the sum of log values from the three central 128bp bins (bin numbers: 447,448,449) for each output track. We performed two types of summarization of the output tracks: 1) directly using the single track that best matched our RNA-seq gene expression data (“Cortex, adult, brain”); 2) using an elastic net model, trained on the predictions from all tracks and all expressed genes (i.e., a matrix of 5,313 tracks-by-13,397 brain expressed genes), to predict population-average gene expression for adult cortex (using GTEx data). As we discuss further in the “sensitivity analysis” section below, the results from these two types of analyses proved similar. Finally, we also performed our evaluation analysis on a smaller set of genes (n=3,401) that overlapped test regions not used to train the Enformer model (Supplementary Fig. 1).

#### Predicting gene expression across individuals

For each individual and each gene, we constructed a personalized DNA sequence input (+/- 196,608 bp) from phased WGS data (separated maternal and paternal DNA sequence inputs were constructed for each individual and each gene). As above, we summed up the log transformed predicted values for the three central 128bp bins (bins numbers: 447,448,449) for each output track. We used two methods to predict final gene expression: 1) used “fine-tuning” as follows: we trained an elastic net model to linearly weight all of Enformer’s 5,313 human output tracks to predict population-average gene expression in the cerebral cortex, using the GTEx RNA-seq data (cortex). Specifically, the elastic net model was fit to predict population average gene expression levels in cortex from the Enformer’s predictions when Reference sequence centered at each gene’s TSS was used as input; 2) we directly selected a single track most representative of cortex gene expression data (“CAGE:brain, adult”). Enformer was used to make separate predictions from the maternal and paternal sequences. For each individual and each gene, we averaged the predictions from the maternal and paternal sequences.

### Statistics and Reproducibility

We used a sample size of 839 (independent subjects) for assessing the significance of the model’s predictions. This sample size is sufficient for assessing significance across individual and per gene, based on previous eQTL analyses^22,23^. We also note that no data from the complete initial dataset (where both WGS and RNA-seq samples passed QC) were excluded from the analyses. Permutation analysis was used to complement the standard false discovery rate (BH) and Bonferroni corrected p-values.

#### Deriving gene-specific Null distribution

Predicted gene expression for the 839 individuals is a function of SNV genotypes for each gene and individual. Thus, we can linearly approximate Enformer’s predictions for each gene and each individual as the weighted sum of the SNVs present in that individual for the given gene^18^. Therefore, to create a null distribution for predictions of gene expression value for each individual and each gene, we assign random attribution weights to each SNV present in the given individual. Specifically, we sample random normally distributed weights for every SNV within the 196,608 bp window around the TSS, and sum them up for each individual genotype as the random gene-specific predictions. For each gene we generate 50 random predictors from which we derive the mean and standard deviation of the absolute Pearson’s correlation to the observed expression values. To assign p-values to Enformer’s correlation to observed gene expression, we use a one-sided T-test and Benjamini-Hochberg procedure to target a 0.05 false discovery rate.

#### Sensitivity analysis

We performed three types of sensitivity analysis, to ensure our cross-individual predictions results are robust. First, we compared the predictions from a single relevant track (CAGE, cortex, adult) and the results when we fine-tuned the predictions with the elastic net model described above (trained on average gene expression prediction from all tracks, using data from GTEx) (**Extended Data Fig. 1A**). Second, we performed model ensembling, whereby we averaged over model predictions on shifted sub-sequences and reverse and forward strands, but this did not impact the sign of significant correlations in ∼96% of cases (**Extended Data Fig. 1B**). Third, when we focused this analysis on 184 genes with known causal SNVs according to previous eQTL analysis^16^, again we observed that while Enformer can make significant predictions, the predicted expression levels are anti-correlated for 80 (43%) of these genes (**Extended Data Fig. 1C, Supplementary Table 1**).

### Training and testing of simple CNN

Our simple CNN was trained on genes that were not located within the regions of Enformer’s test set. During training we used sequences of length 40,001bp from the reference genome centered at the TSS as input to the model and predicted mean log gene expression from the ROSMAP dataset (dorsolateral prefrontal cortex). The length of the input sequence was informed by a recent study^17^. This CNN has a very shallow architecture; it consists of a single convolutional layer with 900 kernels of width 10 and a ReLU activation. We apply a single average pooling layer of size 900 bp that reduces the input of the network to 44 segments. We then apply a single hidden layer of size 200 with ReLU activation before predicting mean gene expression of the given gene. For training we use Mean Squared Error (MSE) loss and Adam optimizer with a learning rate of 0.001 and default hyperparameters. Then, for a random set of 190 individuals, we constructed a maternal and a paternal genomic sequence by inserting all the variant alleles within +/- 20,000bp of the TSS into the Reference sequence. We then made separate predictions for the maternal and paternal sequences and averaged them for every individual. We computed the Pearson’s correlation coefficient between the predicted and observed expression values for these 190 individuals and compared the absolute Pearson’s R to the value that we would expect from our gene specific Null model for variants within +/- 20,000bp of the TSS.

### Driving variant attribution scores using GRAD and ISM

To explore the causes for the negative correlation between Enformer predictions and the observed gene expression values we applied two explainable AI (XAI) techniques on all genes with a significant correlation value (abs(R)>0.2, **Fig. 2A**): ISM and gradients (Grad) ^9,15,24^. Please see the Supplementary Methods for details on rational and methodology, as well as the procedure for identifying the Enformer “driver SNVs” for predictions from WGS data.

### Computing eQTL values and sorting drivers into supported and unsupported drivers

We computed eQTL effect size (ES) for a given SNV as the slope of the linear regression solution that predicts gene expression across individuals from this SNVs genotype, *i.e.* individuals with two copies of the major allele (genotype 0), those with one copy of the major allele (genotype 1) and those with two copies of the minor allele (genotype 2). The slope of the regression with the genotype of each SNV represents how much expression changes with an additional copy of the minor allele. Positive or negative slopes determine the direction of SNV effect on gene expression. Based on the eQTL ES and ISM attribution values for each SNV, one can distinguish between supported and unsupported drivers. Supported drivers’ attributions have the same sign as the eQTL ES and unsupported drivers have the opposite sign.

### Data Availability

Genotype, RNA-seq, and DNAm data for the Religious Orders Study and Rush Memory and Aging Project (ROSMAP) samples are available from the Synapse AMP-AD Data Portal (**Accession Code: syn2580853**) as well as RADC Research Resource Sharing Hub at www.radc.rush.edu.

### Code Availability

Scripts for running the analyses presented, as well as intermediate results are available from: https://github.com/mostafavilabuw/EnformerAssessment^25^

---

**Extended Figure 1.**
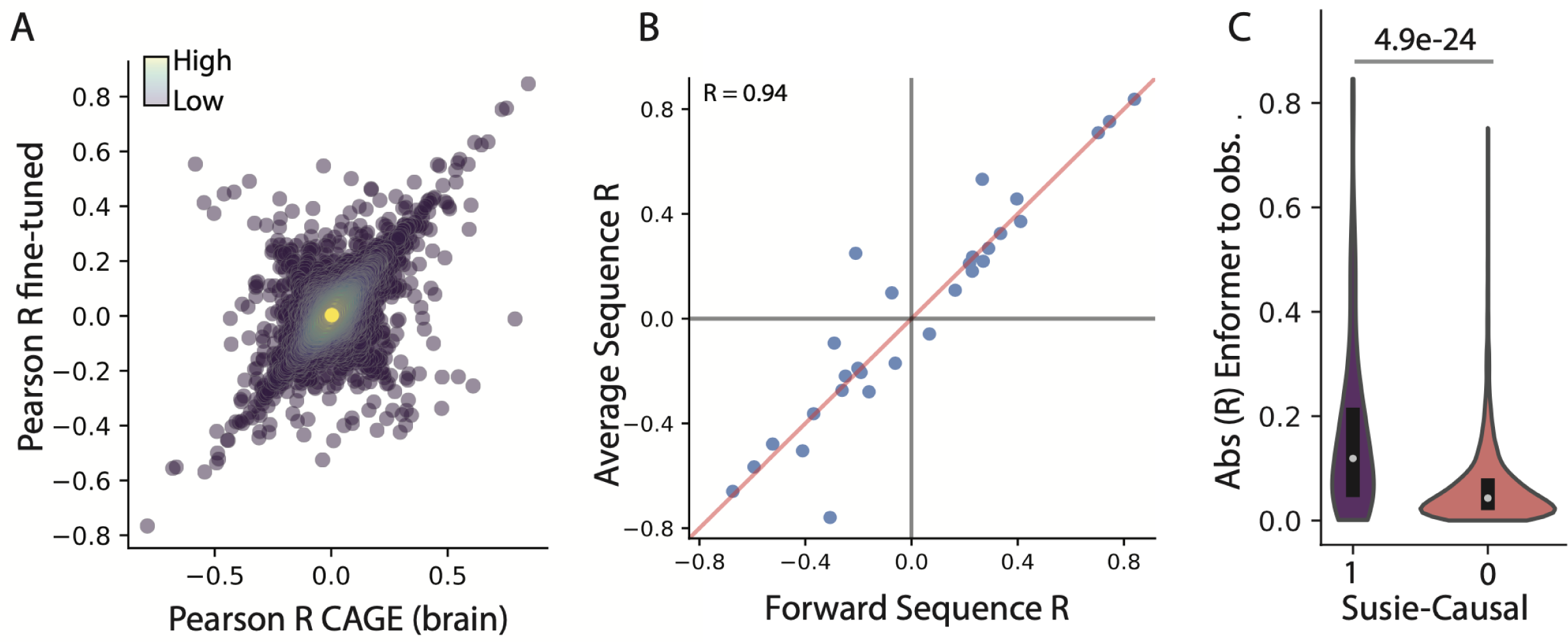
Sensitivity analysis for Enformer predictions. (A) Density plot, where each dot represents a gene (n=13,397). X-axis shows Pearson R coefficients for Enformer predictions for the single most relevant track (“CAGE,brain,adult”) and y-axis shows the fine-tuned cortex model from all human tracks. Color depicts local density. (B) Pearson R coefficients across 839 individuals between observed expression and the predicted CAGE track from a single forward-stranded input sequence centered at the TSS (x-axis) versus the average over forward-stranded sequences which were shifted by -3, -2, -1, 0, 1, 2, 3 bp, and a reverse-stranded input sequence centered at the TSS (y-axis). Data shown for a random subset of loci (n=30). Orange line: diagonal line where x and y-axis have the same value. The correlation coefficient between values on x-axis and y-axis is R=0.94 (C) Absolute Pearson R coefficients between Enformer predictions and observed gene expression for sets of genes with one causal SNP and all others. Causal genes determined by the Susie algorithm (“Susie-Causal”). Edges of the box indicate the 25th and 75th percentiles, and the central mark indicates the median (N1=183 genes fine-mapped with Susie, N2= 6625 genes without fine-mapped variants, two-sided Wilcoxon rank-sum test, for each gene R coefficient computed using n=839 individuals).

**Extended Figure 2.**
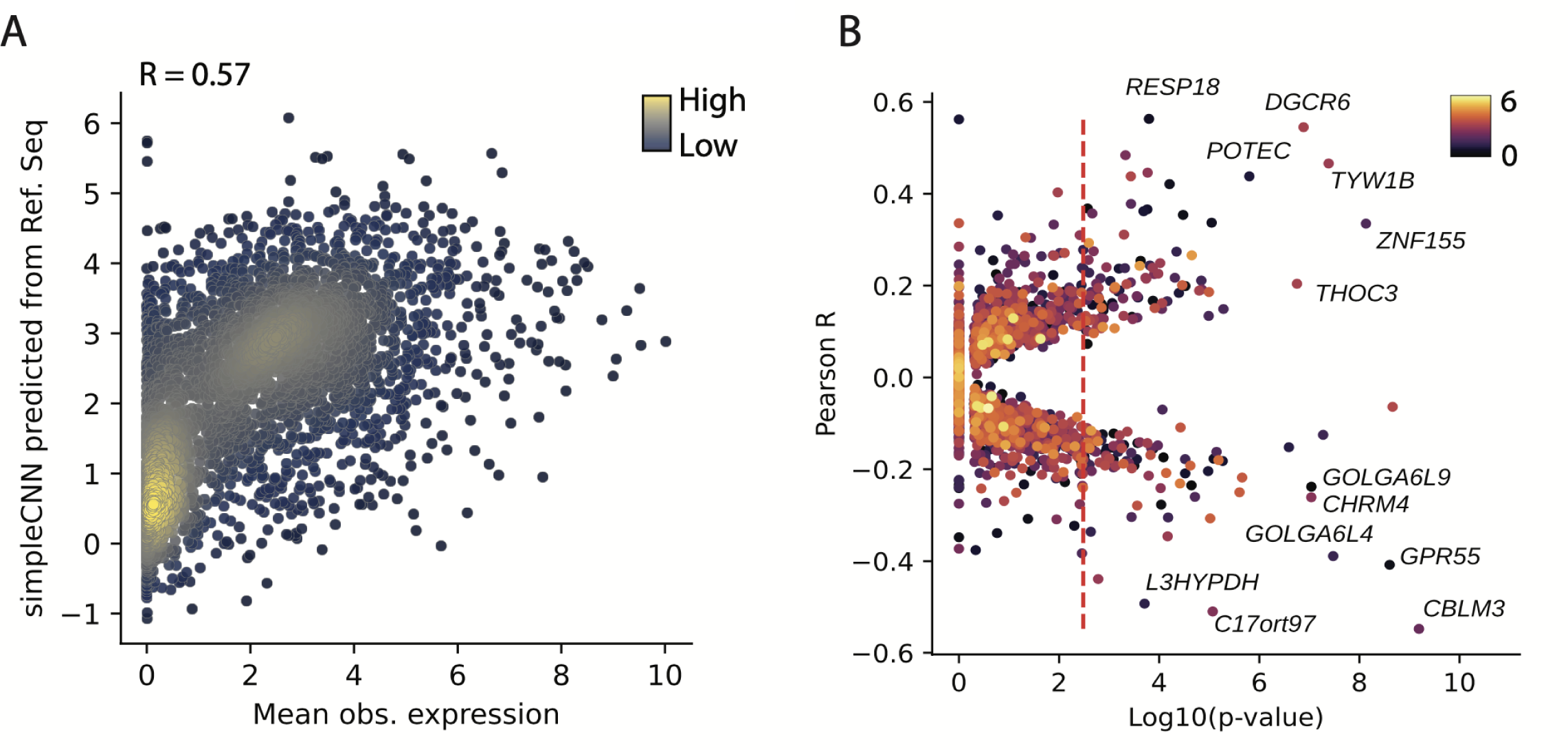
Performance of the simple CNN model. (A) Density plot of observed population-average expression of test set genes (n=3,401 genes) in cerebral cortex versus simple CNN’s predicted gene expression from the Reference sequences. This plot only displays genes which could be assigned to Enformer’s test set. Colors depict local density. (B) Y-axis shows Pearson R coefficients between observed expression values and a simple CNN’s predicted values per individual. X-axis shows the negative log10 p-value computed with a gene-specific Null model (one-sided T-test, n=50 independent samples per gene; Supplementary Method). The color represents the predicted mean expression. Red dashed line indicates FDRBH=0.05.

## Supplementary Methods

### S1. Data Pre-processing

#### RNA-seq gene expression data from the ROSMAP cohort

RNA-seq data were collected from dorsolateral prefrontal cortex (DLPFC) tissue of 1118 individuals^1^, and preprocessed with the pipeline described by Felsky and colleagues^3^. In brief, we applied TMM normalization (using edgeR calcNormFactors) to the raw counts to estimate the effective library size of each individual. We then applied voom/limma to regress out confounds and convert the counts into log_2_(CPM). Technical covariates included:

1. batch, study (ROS or MAP)
2. RNA integrity number, postmortem interval
3. Library size
4. Log PF number of aligned reads
5. PCT_CODING_BASES
6. PCT_INTERGENIC_BASES
7. PCT_PF_READS_ALIGNED
8. PCT_RIBOSOMAL_BASES
9. PCT_UTR_BASES
10. PERCENT_DUPLICATION
11. MEDIAN_3PRIME_BIAS
12. MEDIAN_5PRIME_TO_3PRIME_BIAS
13. MEDIAN_CV_COVERAGE.

Biological covariates, including 1) age, 2) sex, and 3) top 10 expression principal components. Both biological and technical covariates were regressed out from log raw read counts. Only genes with mean log_2_(CPM) > 2 were included. Mean expression values were retained for downstream analysis.

#### WGS data from the ROSMAP cohort

The variant call files for whole genome sequencing (WGS) data from the ROSMAP in variant call format (VCF) were obtained from the Synapse repository (syn117074200). The coordinates of variant calls (GRCh37) were converted to GRCh38 coordinates using the Picard LiftoverVcf tool (http://broadinstitute.github.io/picard). The Eagle software2 version 2.4.1 was used to phase the genotypes with the default setting.

### S2. Driving variant attribution scores

To explore the causes for the negative correlation between Enformer predictions and the observed gene expression values we applied two explainable AI (XAI) techniques on all genes with a significant correlation value (abs(R)>0.2, **Fig. 2A**): ISM and gradients (Grad) ^3–5^. Below, we describe both methods, however, we only present the result of the ISM method in the main text, due to its superior performance. Specifically, while there was a moderate correlation between attributions computed with Grad and ISM (mean *Pearson R* = 0.28, **Fig. S5**), we found that linear decomposition with ISM generated a better approximation of Enformer’s predictions (**Fig. S6**), and was able to accurately approximate Enformer’s predictions for 95% of the examined genes (R>0.2, p<10^-8^).

#### Linear approximation of NN models

To understand the impact of each variant to Enformer’s prediction of gene expression for 839 individual sequences we linearly approximate the neural network function using the Taylor expansion. In contrast to Deep Neural Networks (DNNs), linear functions are easy to interpret. To approximate the value of a complex function with arbitrary input *s*, the Taylor approximation linearly decomposes the function value at *s* into the value of a function at nearby position *f(s_0_)* and the derivative of the function at *s_0_* multiplied by the difference between the position of interest *s* and s*_0_*^6^:

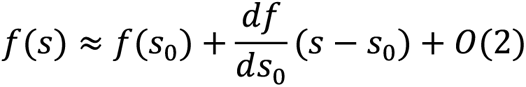

Where O(2) represents the second order term that is truncated in the linear approximation. In our problem, the function values at Reference genotypes are *f(s_0_)* and the gradient with respect to the Reference genotype for the selected output track give 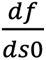. The positions of interest are the individual genome sequences that contain different sets of variants *L_i_*. In a one-hot encoded input, to get the difference from the Reference sequence to the individual genomic sequence, for each loci, we have to delete the Reference base (set b_0_ from 1 to 0) and introduce a new base (set b_1_ from 0 to 1). If we assume that these changes in each set of variants *L_i_* are independent and linear, we can approximate the function value for an individual’s genomic sequence *s_i_* as the sum of changes from the reference sequence times the gradient at these positions.

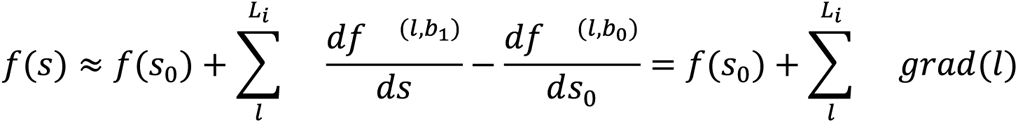

The gradients, denoted *grad(l)*, are computed for “free” in the forward pass with the Reference sequence and can therefore be used directly. The gradient at the Reference represents a local numerical approximation.

Another approximation of the gradient is defined by finite differences from the reference to a sequence with a single variant, which is also referred to as *in-silico mutagenesis* (ISM).

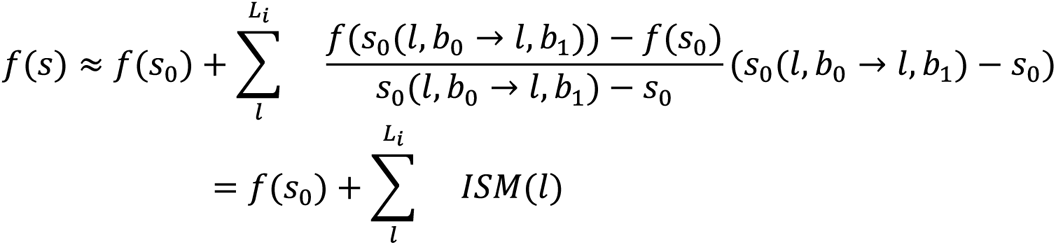

We note that linear approximation is not a perfect approximation of the non-linear neural network. However, the linear weights generally cover the most impactful contributions to the predictions and we only derive our explanations for genes for which the linear approximation correlates significantly with Enformer’s predictions.

#### Computation of Gradients

To compute the gradient at the Reference sequence we performed forward passes with all genomic sequences from the Reference genome of length 196,008b centered at the TSS of each gene. As a predictor for gene expression we selected track_idx = 4980 (“CAGE:brain, adult”) and summed over the central windows 447, 448, 449 to compute the gradient for the predicted gene expression. The gradient attribution at each loci *l* is then computed as the difference between gradients at variant base *b_1_* and the base in the reference sequence *b_0_* (see Taylor expansion above).

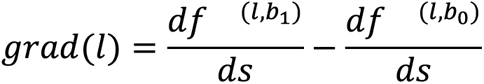

#### Computation of ISM

To compute ISM at the Reference sequence we performed forward passes with all genomic sequences from the Reference genome of length 196,008b centered at the TSS. As a predictor for gene expression we selected track_idx = 4980 (“CAGE:brain, adult”) and summed over the central windows 447, 448, 449. We then separately inserted the most common alternate allele at each possible SNV position within the 196Kb window into the Reference sequence, setting the reference base b0 at the loci l to zero and the new b1 base to one. We consider possible SNV positions to be any position where at least one subject in the dataset has a SNV. For each of these sequences with a single base change from the Reference, we performed a forward pass and computed the predicted gene expression as the sum of the three windows. Consequently, to get all ISM attributions for each gene we had to perform as many forward passes as variants are around the gene. The finite difference for each variant is the difference between the predicted value from the variant sequence and the reference sequence.

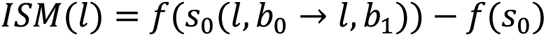

#### Selection of drivers

To contribute to the variance of predictions across different individual genomes, a variant has to have a large attribution but also it has to occur in the right set of individuals, that is the genotype has to be correlated with gene expression. To determine which variant’s weight contributes most to the linear approximation, i.e. sum over linear weights (ISM) times genotype, we sorted the ISM attributions (i.e., SNV weights) in order of their absolute value with the largest attribution at first. We then iteratively included these attributions to the sum and determined which attributions improved the correlation of the sum with the original Enformer predictions (CAGE,adult,brain). SNVs were defined as drivers if adding their attribution to the partial sum increased the correlation with the Enformer prediction by at least 5% of the correlation between Enformer predictions and sum over weights from all variants and the variants’ genotype times attribution also significantly correlated with the Enformer predictions themselves (i.e., p<0.01, Bonferroni correction).

### S3. Defining and extracting sequence motifs around drivers

To determine if specific sequences are overrepresented around driver SNVs, we extracted 13-mer sequences (*i.e.* 7-mers on each side including the driver) around every driver and clustered them into motifs. For each pair of 13-mers we found their best alignment without introducing gaps, requiring that they had at least four aligned bases, and that the location of both SNVs would be included in the aligned part. We scored the alignments by the number of aligned bases and divided this score by five to determine the fraction of bases out of five aligned bases. We then performed agglomerative clustering with complete linkage and a threshold of 0.8 meaning that at least four bases had to be aligned between all 13-mers in a cluster. This procedure produced 256 sequence clusters from the 13-mers around 1091 drivers in the reference sequence of 251 genes (**Fig. S8A**). The 13-mers of the variant sequence of these 1091 drivers clustered into 261 sequence patterns (**Fig. S8B**), and the reference sequence at the main drivers of these 251 genes clustered into 84 sequence patterns (**Fig. S8C**). We then determined the enrichment of driver types (i.e. supported or unsupported) for each cluster. We used Fisher’s exact test to compute the p-value of this enrichment, and Benjamini-Hochberg procedure to correct for multiple testing. None of the clusters were significant when targeting a FDR of 0.05.

### S4. Defining and extracting significant attributional motifs from ISM

Since sequence-based models learn sequence motifs that are predictive of functional properties like gene expression, we reasoned that examining “*attributional motifs”* will be informative in understanding when predictions go wrong. We thus defined attributional motifs as short regions of DNA where a few consecutive bases all receive a reasonably high feature attribution (defined below). Absence of such attributional motifs in a given genomic loci indicates insufficient training data for learning the regulatory logic that enable coherent predictions.

Specifically, we computed ISMs for 41bp around all driver SNVs at the Reference sequence and the variant sequence (Reference with driver SNV inserted). We also computed ISMs within windows of 2000bp around each gene’s TSS (**Fig. S11A**). To compare effect sizes of SNVs across genes, we standardize their effect by the maximum absolute ISM in 2000bp around the TSS, so that each ISM values around the drivers represent the percentage or fraction of the maximum ISM value that we observe (**Fig. S11B,-C**). This way, we determine how important a base change is compared to effects of other bases along the entire gene, and we can ignore potential motifs with negligible impact on Enformer’s predictions. We use the standardized ISMs around the drivers to determine in the reference and also the variant sequence how many subsequent base changes have ISM values with more than 5%, 10%, 20% and 50% of the maximum absolute ISM (Supplementary Table 3), and if they don’t surpass these thresholds, we determine the distance to the nearest base above these thresholds (Fig S14D, for 10% threshold).

## Supplementary Figures

**Figure S1.**
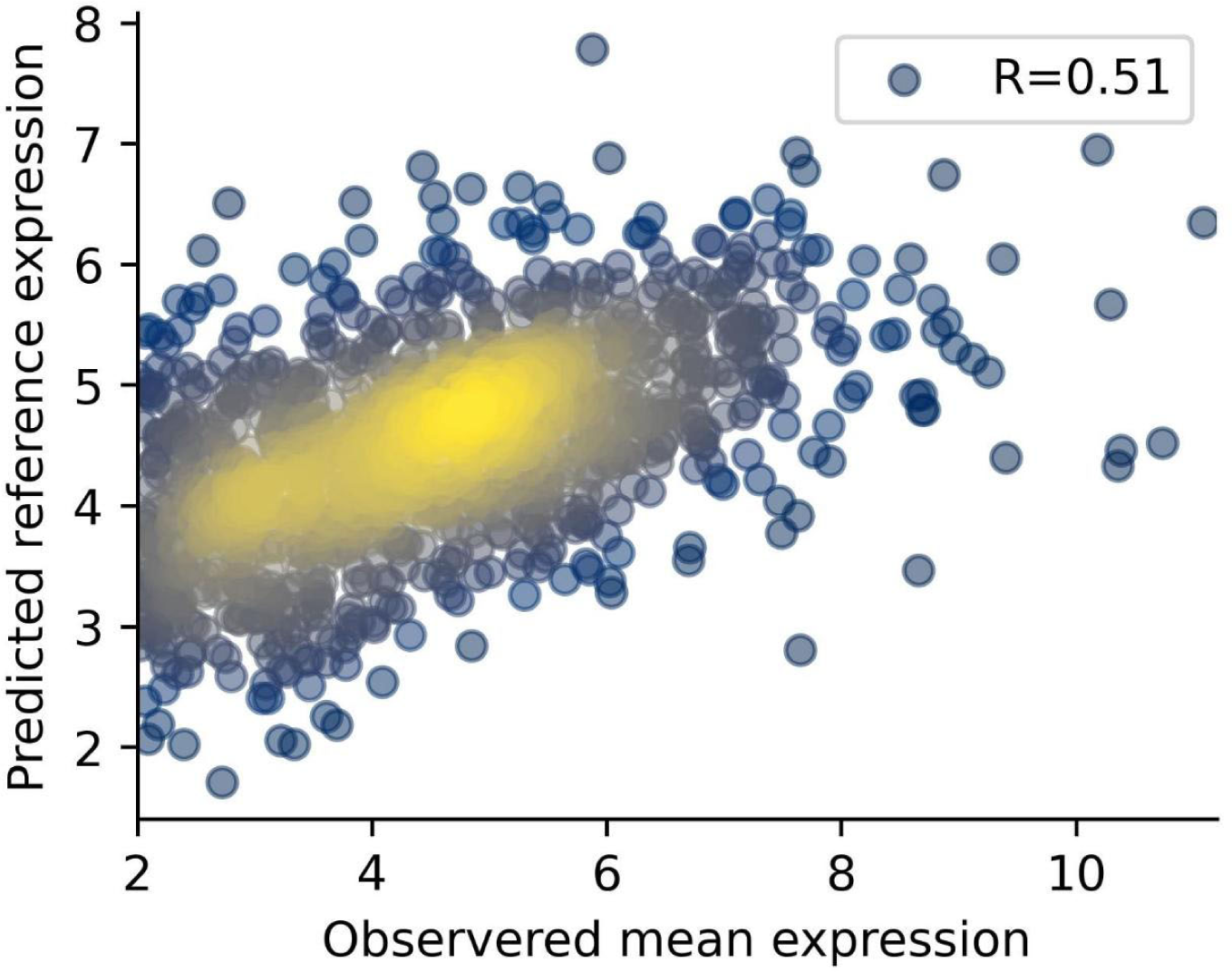
Observed population-average expression from cerebral cortex versus Enformer’s predicted gene expression using the fine-tuned model. Inputs are Reference sequences centered at TSS of each tested gene. This plot only displays genes which could be assigned to Enformer’s test set.

**Figure S2.**
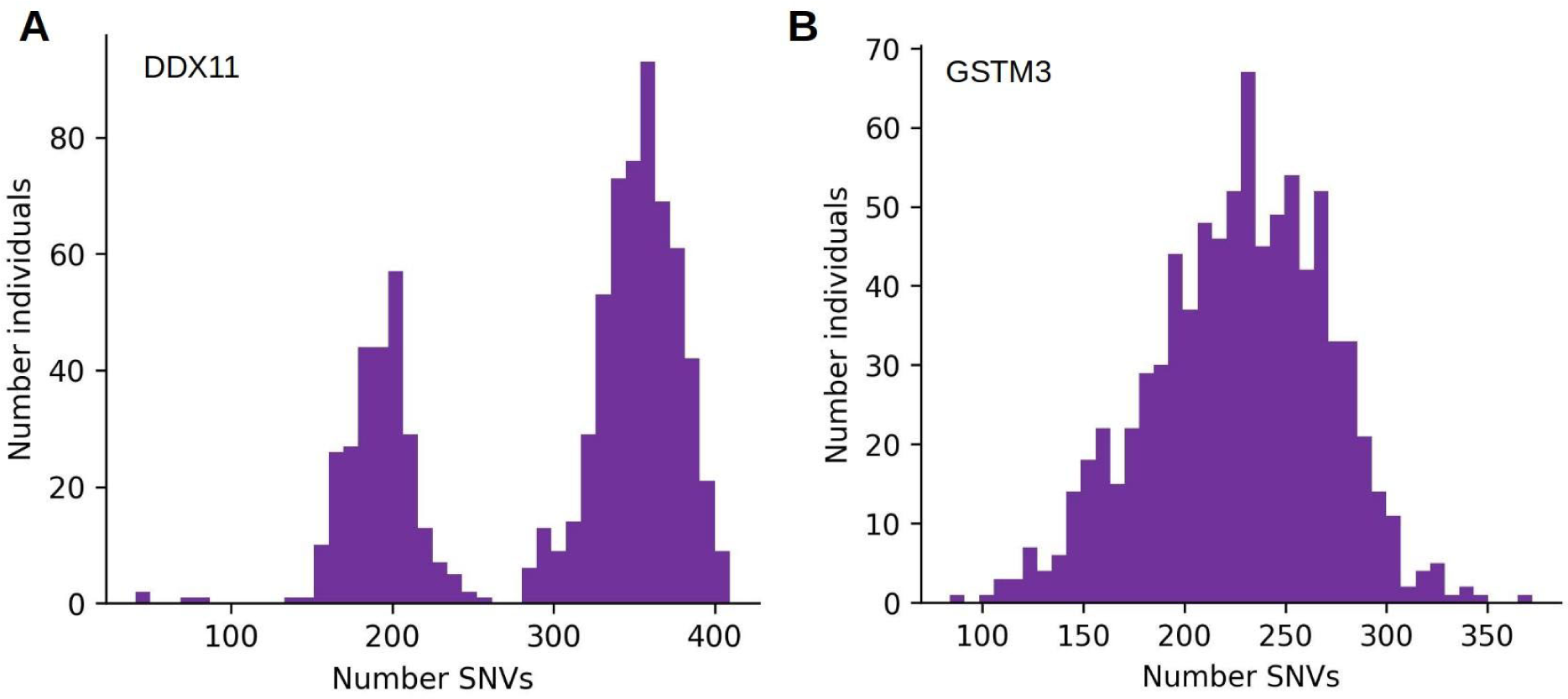
Distribution of the number of SNVs per individual for genes *DDX11* and *GSTM3*.

**Figure S3.**
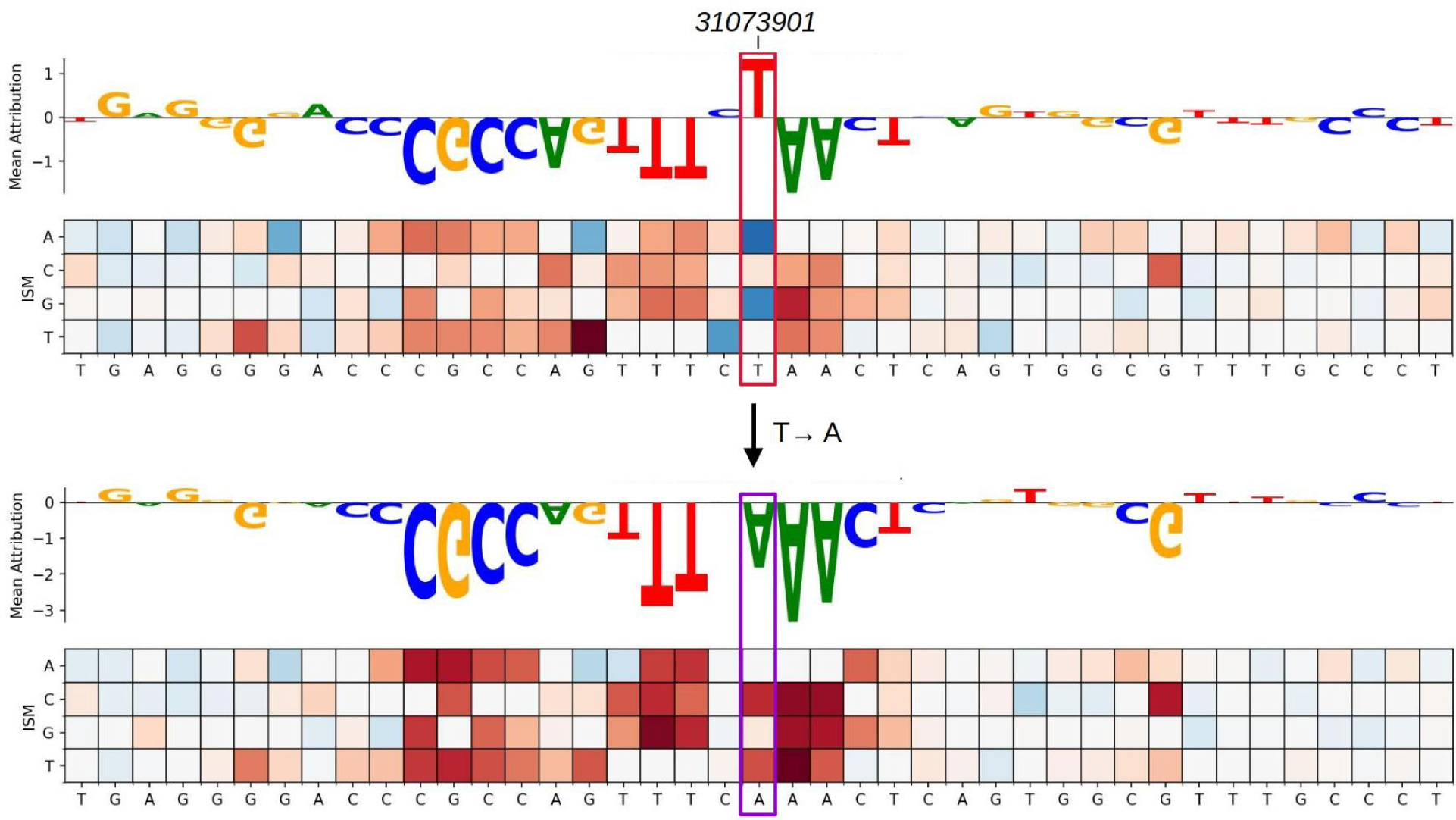
*In silico* mutagenisis for the strongest variant in *DDX11* at rsID rs7953706 (located at chr 12, 31073901). Top) top plot shows the mean attributions for the Reference sequence from ISM and the heat map below shows the individual ISM values for each of the three variants. Red indicates that replacing the base at this position with the variant allele would increase expression and blue indicates decreasing expression. Bottom) Mean attributions for the Reference sequence with the variant rs7953706 inserted and below the individual ISM values for each of the three possible variants.

**Figure S4.**
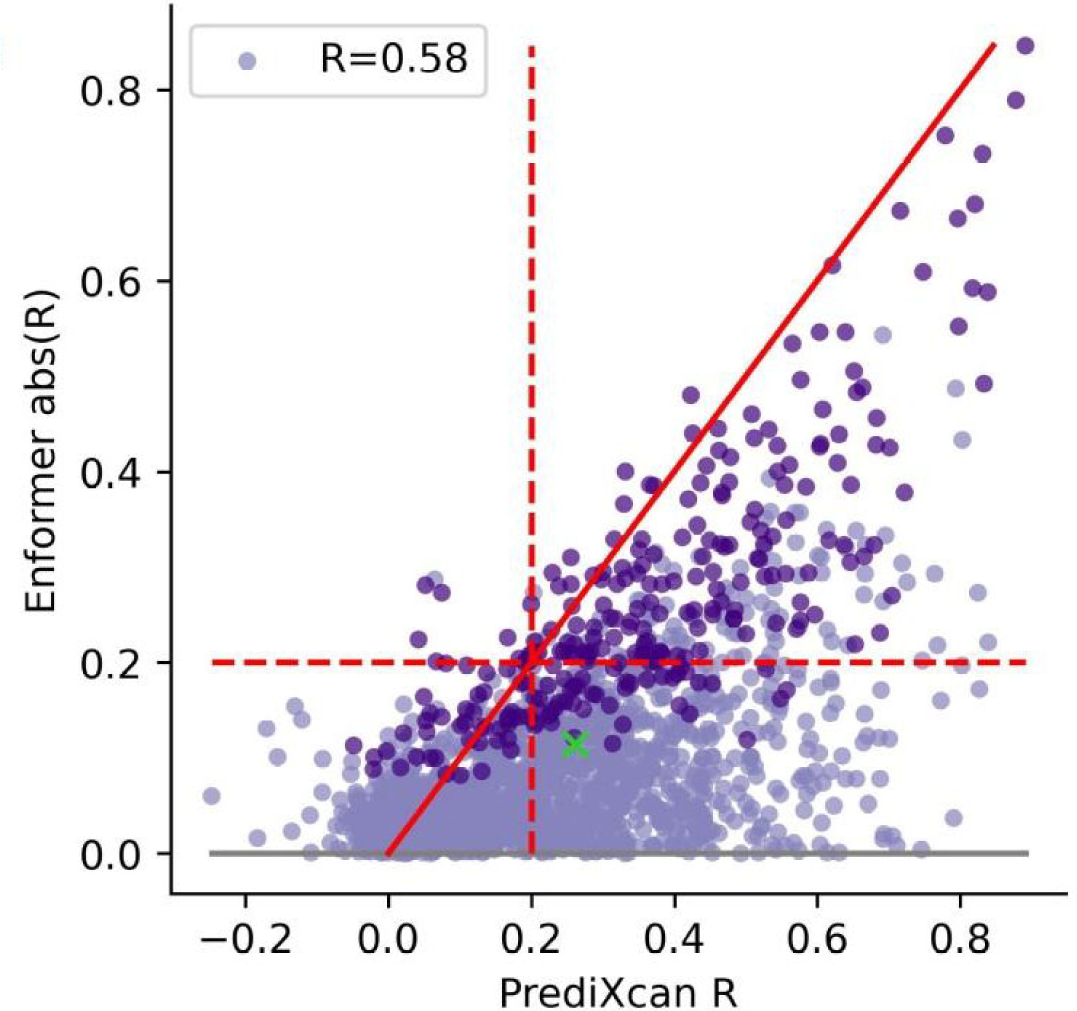
Absolute Pearson Rs between PrediXcan predicted versus observed expression across 839 individual genomic sequences (x-axis) versus Enformer’s absolute Pearson R values (y-axis). Red lines indicate threshold for significance of positive predictions (R>0.2, FDR_BH_<0.05). Green cross represents the location of the mean across all x- and y-values. Darker colored dots represent genes that possess significant correlation compared to gene’s SNV-specific distribution as shown in Figure 2A (FDR_BH_<0.05).

**Figure S5.**
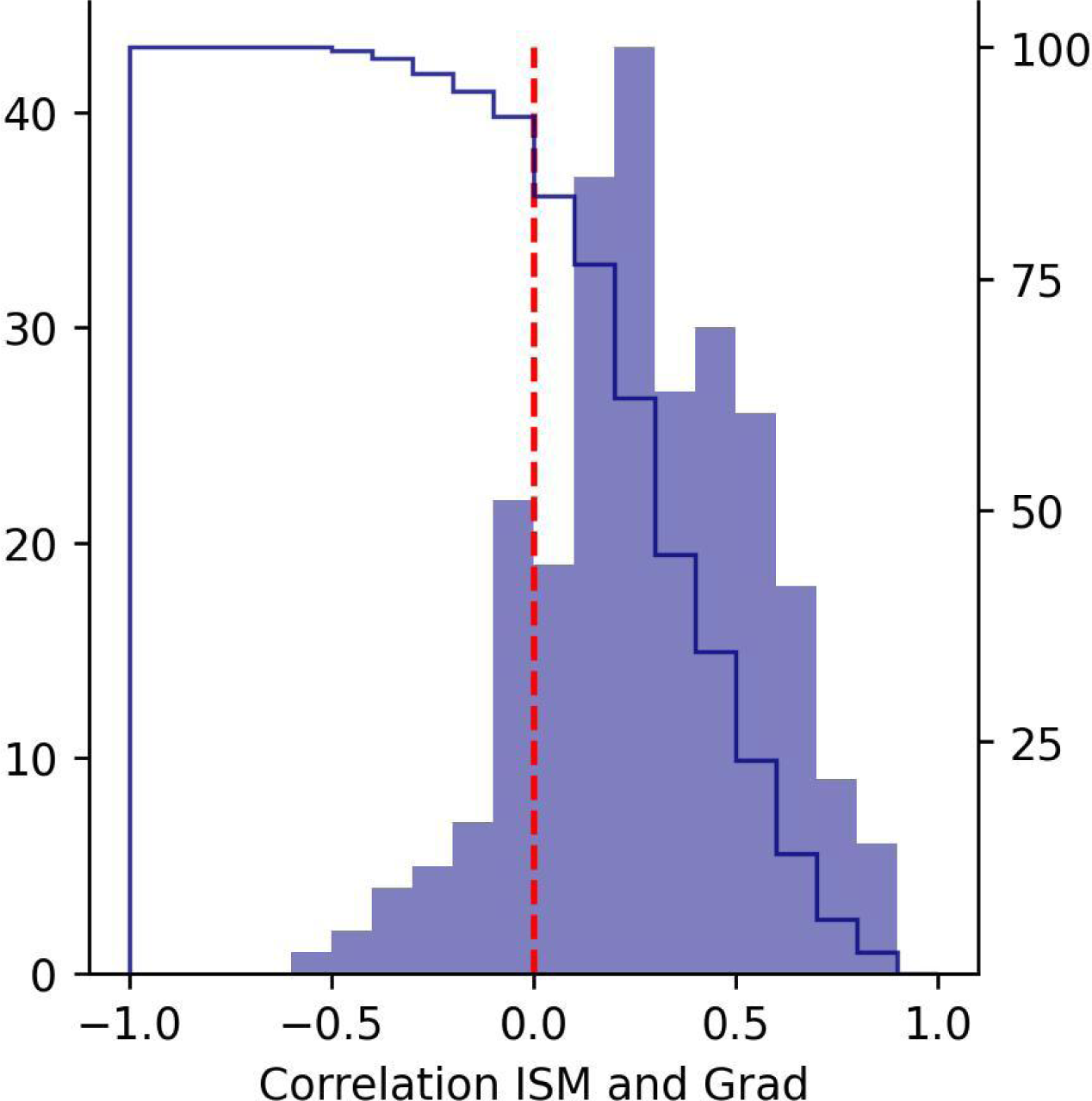
Correlations between ISM and Grad attributions for all variants of every gene. Distribution (blue bars, left axis) and reverse cumulative percentage (dark blue line, right axis)

**Figure S6.**
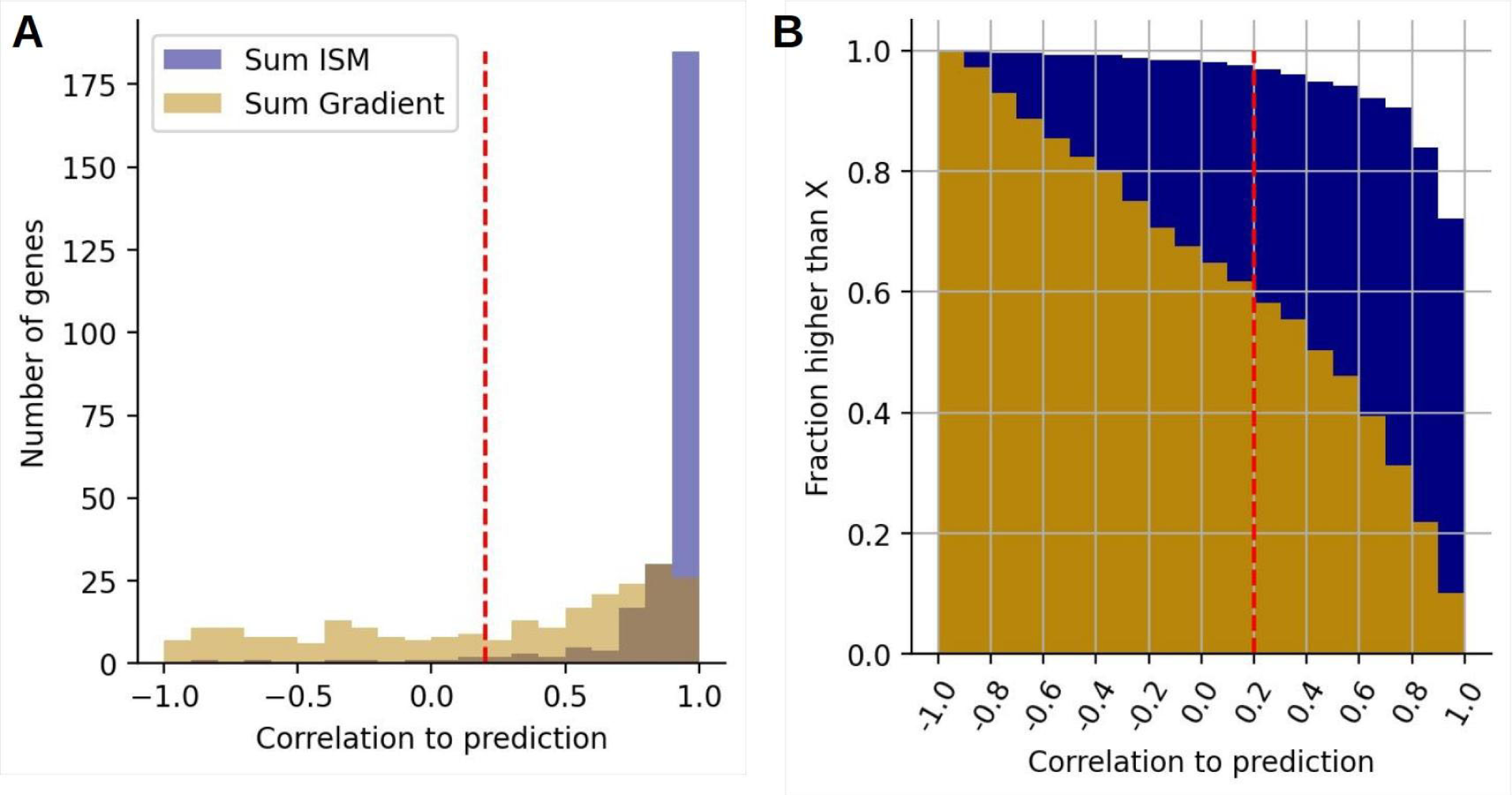
Correlations between sum of attributions (linear approximation) and Enformer predictions (CAGE,adult,brain track) for Grad (golden) and ISM (navy blue). A) Histogram of correlations B) Faction of tested genes with a correlation higher than indicated on the x-axis (sum from right to left).

**Figure S7.**
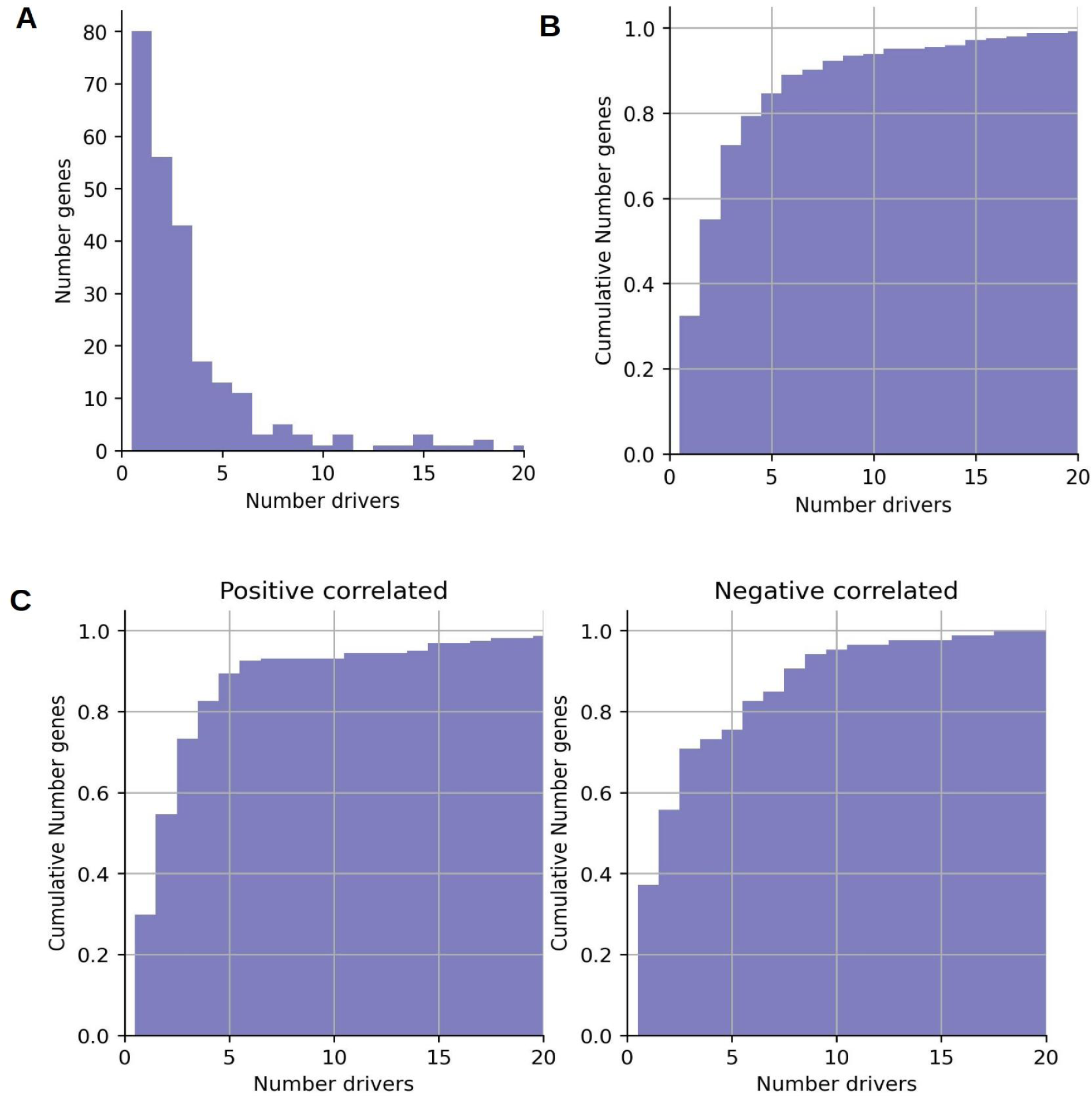
A) Number of genes with number of SNV drivers selected by forward method (see Methods). B) Fraction of genes with at most X SNV drivers. C,D) Fractions of genes with at most X SNV drivers for genes with positively correlated predictions to observed expression and negatively correlated predictions.

**Figure S8.**
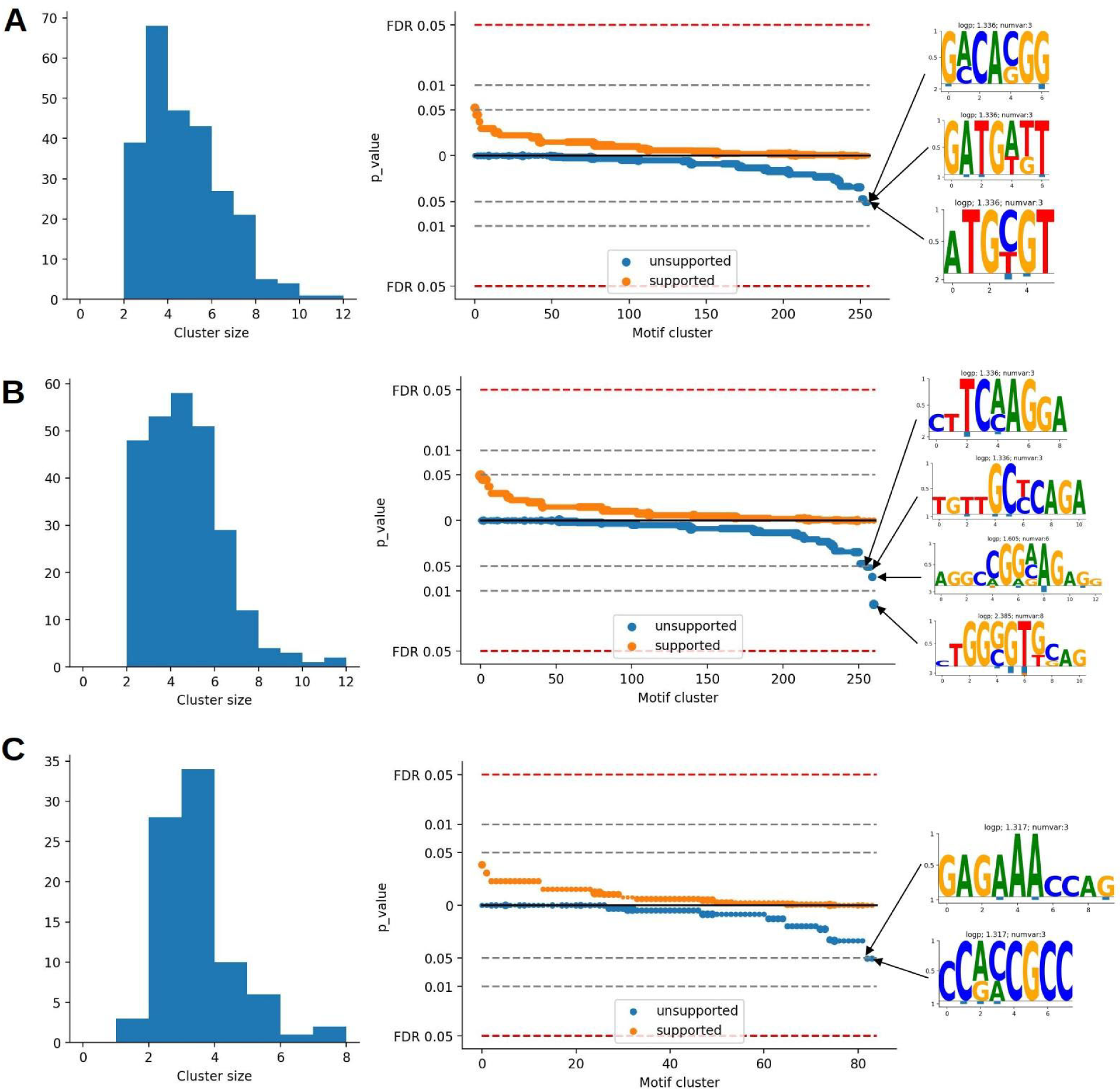
Clustering 13-mers around SNV drivers at the reference and variant sequences to determine motifs that are enriched for supported and unsupported drivers. A) 13-mer clusters around all SNV drivers with the Reference base at driver SNV. B) 13-mer clusters with the variant base at driver loci. C) 13-mer clusters with the Reference base at the main driver SNVs.

**Figure S9.**
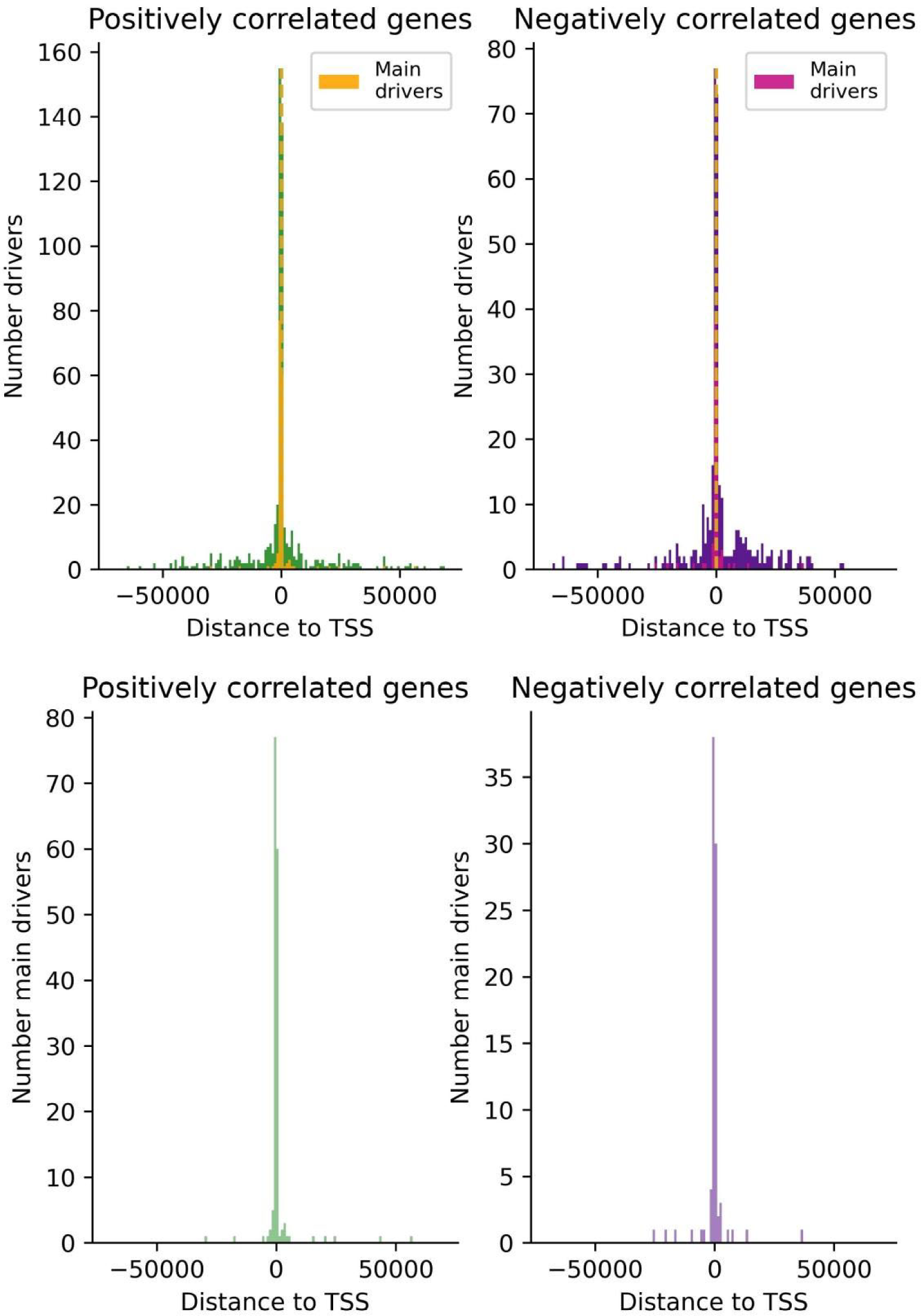
Top: Location of driver SNVs along the entire input sequence within 1000 bp windows. Bottom: Main driver locations along the entire sequence within 1000 bp windows

**Figure S10.**
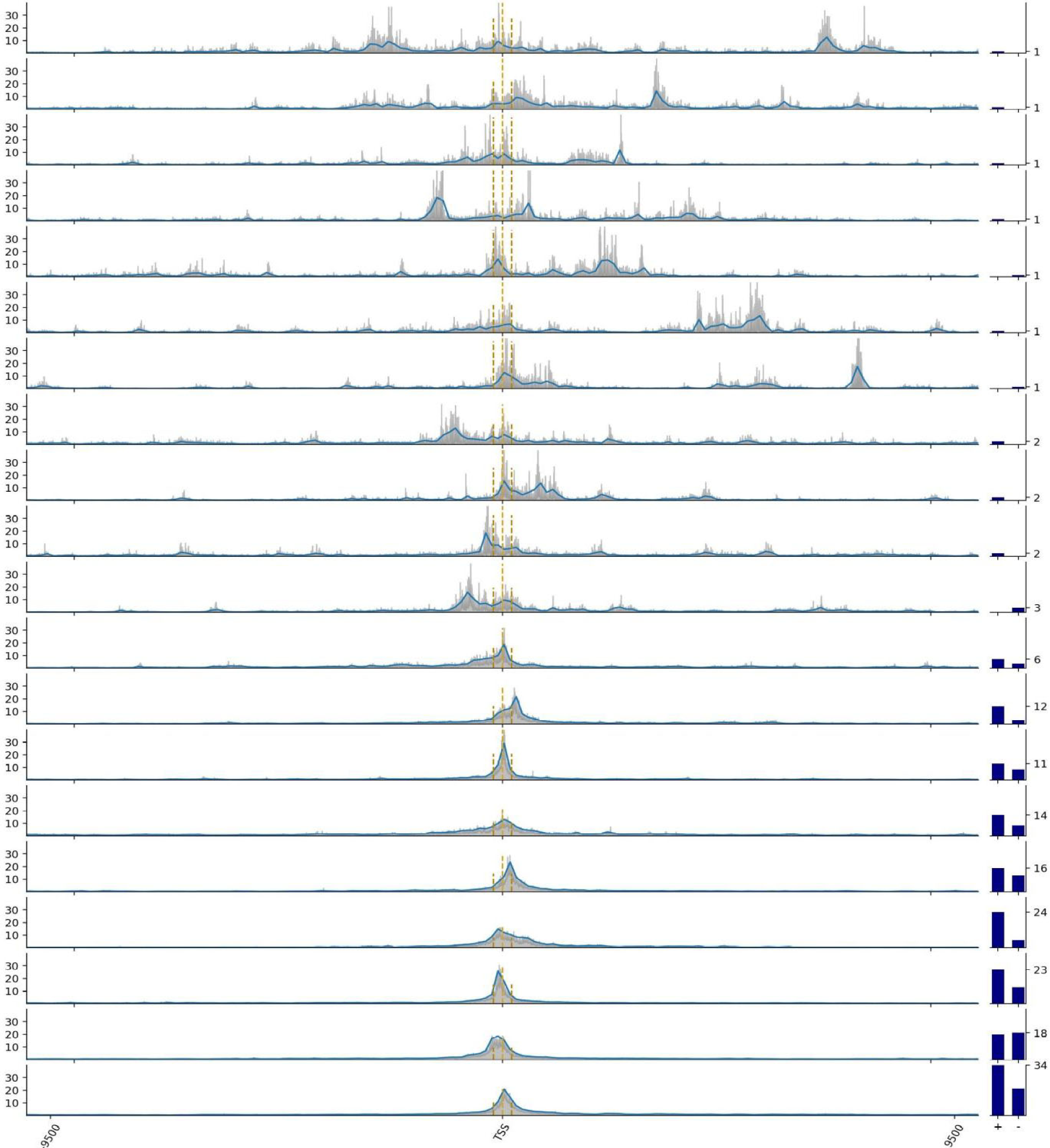
Gradient attribution clusters from clustering standard deviation of Grad attributions within 128bp. Gradient attributions of 251 genes were clustered with Euclidean distance on the standard deviation of standardized attributions within windows of 128 bp along the entire 196,008bp input sequence. The figure shows a 20,000 bp window around the TSS with the mean of the absolute standardized attribution of 20 clusters that were found by agglomerative clustering with complete linkage. Grey bars show the mean absolute standardized attribution for each cluster and blue line shows the mean standard deviation at this position. Right bars show how many positive (+) and negative (-) correlating genes are part of these clusters. Vertical golden dashed line in the center indicates the location of TSS, while dashed lines around it indicate the boundaries of the three central windows of Enformer output tracks from which these gradients were derived from.

**Figure S11.**
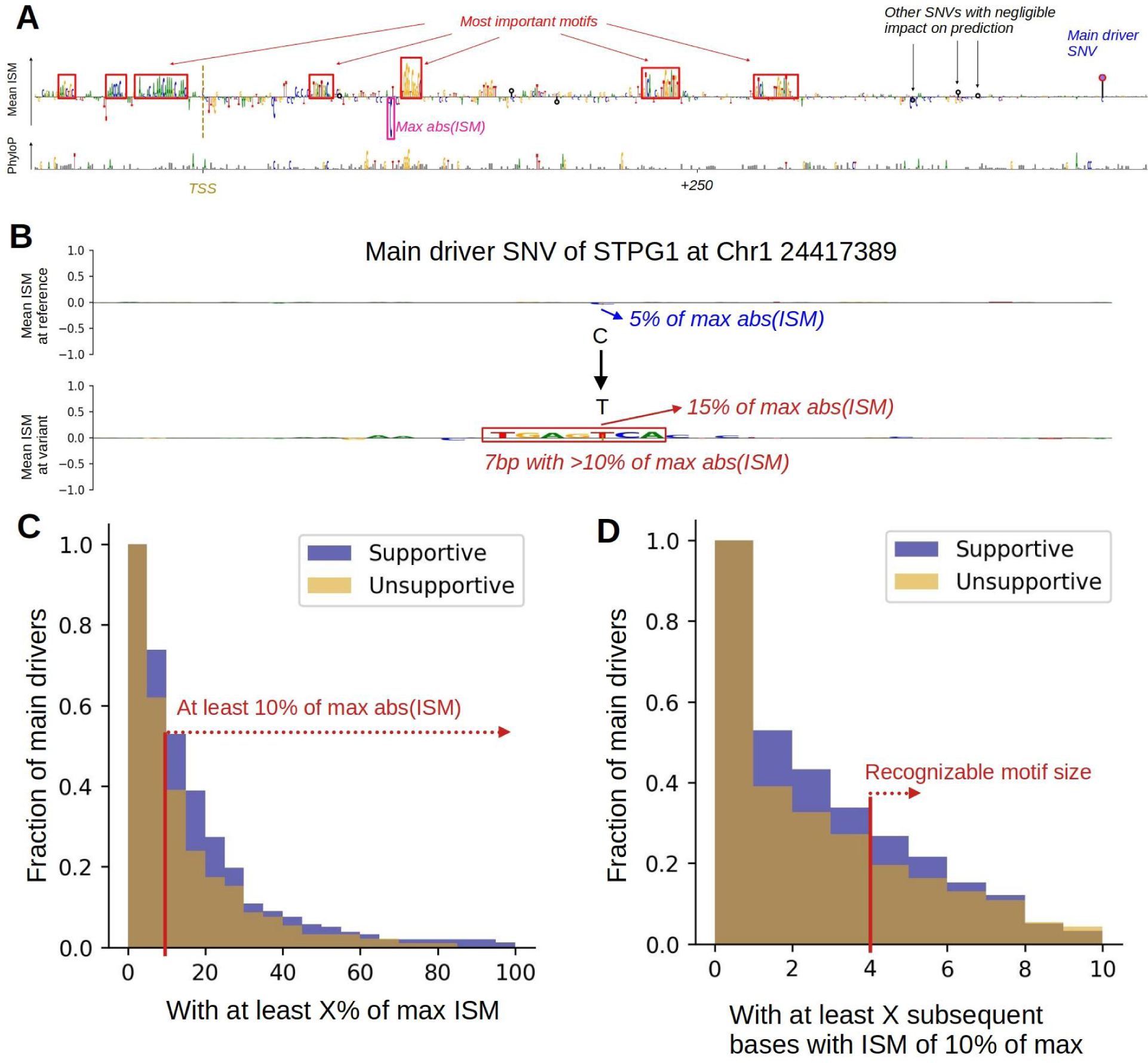
ISM effect of main drivers compared to largest observed absolute ISM value in 2000 bp window around TSS. A) Top: Mean ISM effect of changing one base pair to another one near the TSS of *STPG1* gene. Bottom: PhyloP score for each base pair. PhyloP scores above 1.3 are shown as letters, others as grey bars. Main recognizable motifs are highlighted in red frames. Common SNVs are shown as dots, color indicates Pearon’s R of SNVs genotype to observed expression, red is positive, white is near zero, and blue is negative correlation, location of dot indicates the ISM for changing the reference to the variant base. B) Fraction of mean ISM effect to maximum absolute effect found in 2000bp window around the TSS (see A) for 40bp around the main driver of *STPG1* gene. ISM values are shown in the Reference sequence (top) and in the variant sequence (bottom, reference sequence with the inserted variant). Changing bases from C to T at chr1-24417389 introduces a repressive t motif which has more than 10% the effect of the maximum absolute ISM. C) Fraction of main drivers (supported:indigo, unsupported:gold) for 251 genes with at least X% of maximum absolute ISM. 40% of unsupported and 52% of supported drivers have an effect of 10% or more on Enformer’s predictions. D) Fraction of main drivers that exert 10% of the maximum ISM effect and are surrounded by X bases with ISM effect that also exert more than 10% of the maximum absolute ISM effect.

## References

1. Avsec, Ž., et al. Effective gene expression prediction from sequence by integrating long-range interactions. Nat. Methods 18, 1196–1203 (2021).

2. Avsec, Ž., et al. Base-resolution models of transcription-factor binding reveal soft motif syntax. Nat. Genet. 53, 354–366 (2021).

3. Eraslan, G., Avsec, Ž., Gagneur, J. & Theis, F. J. Deep learning: new computational modelling techniques for genomics. Nat. Rev. Genet. 20, 389–403 (2019).

4. Zhou, J., et al. Whole-genome deep-learning analysis identifies contribution of noncoding mutations to autism risk. Nat. Genet. 51, 973–980 (2019).

5. Zhou, J. Sequence-based modeling of three-dimensional genome architecture from kilobase to chromosome scale. Nat. Genet. 54, 725–734 (2022).

6. Park, C. Y., et al. Genome-wide landscape of RNA-binding protein target site dysregulation reveals a major impact on psychiatric disorder risk. Nat. Genet. 53, 166–173 (2021).

7. De Jager, P. L., et al. A multi-omic atlas of the human frontal cortex for aging and Alzheimer’s disease research. Sci Data 5, 180142 (2018).

8. Kelley, D. R., Snoek, J. & Rinn, J. L. Basset: learning the regulatory code of the accessible genome with deep convolutional neural networks. Genome Res. 26, 990–999 (2016).

9. Zhou, J. & Troyanskaya, O. G. Predicting effects of noncoding variants with deep learning-based sequence model. Nat. Methods 12, 931–934 (2015).

10. Yuan, H. & Kelley, D. R. scBasset: sequence-based modeling of single-cell ATAC-seq using convolutional neural networks. Nat. Methods (2022) doi:10.1038/s41592-022-01562-8.

11. Maslova, A., et al. Deep learning of immune cell differentiation. Proc. Natl. Acad. Sci. U. S. A. 117, 25655–25666 (2020).

12. Chen, K. M., Wong, A. K., Troyanskaya, O. G. & Zhou, J. A sequence-based global map of regulatory activity for deciphering human genetics. Nat. Genet. 54, 940–949 (2022).

13. Kim, D. S., et al. The dynamic, combinatorial cis-regulatory lexicon of epidermal differentiation. Nat. Genet. (2021) doi:10.1038/s41588-021-00947-3.

14. Zhou, J., et al. Deep learning sequence-based ab initio prediction of variant effects on expression and disease risk. Nat. Genet. 50, 1171–1179 (2018).

15. Novakovsky, G., Dexter, N., Libbrecht, M. W., Wasserman, W. W. & Mostafavi, S. Obtaining genetics insights from deep learning via explainable artificial intelligence. Nat. Rev. Genet. (2022) doi:10.1038/s41576-022-00532-2.

16. Wang, Q. S., et al. Leveraging supervised learning for functionally informed fine-mapping of cis-eQTLs identifies an additional 20,913 putative causal eQTLs. Nat. Commun. 12, 3394 (2021).

17. Karollus, A., Mauermeier, T. & Gagneur, J. Current sequence-based models capture gene expression determinants in promoters but mostly ignore distal enhancers. Genome Biol. 24, 56 (2023).

18. Gamazon, E. R., et al. A gene-based association method for mapping traits using reference transcriptome data. Nat. Genet. 47, 1091–1098 (2015).

19. Reshef, Y. A., et al. Detecting genome-wide directional effects of transcription factor binding on polygenic disease risk. Nat. Genet. 50, 1483–1493 (2018).

## Methods References

20. Bennett, D. A. et al. Religious Orders Study and Rush Memory and Aging Project. J. Alzheimers. Dis. 64, S161–S189 (2018).

21. Mostafavi, S., et al. A molecular network of the aging human brain provides insights into the pathology and cognitive decline of Alzheimer’s disease. Nat. Neurosci. 21, 811–819 (2018).

22. Battle, A., et al. Characterizing the genetic basis of transcriptome diversity through RNA-sequencing of 922 individuals. Genome Res. 24, 14–24 (2014).

23. GTEx Consortium. Genetic effects on gene expression across human tissues. Nature 550, 204–213 (2017).

24. Sundararajan, M., Taly, A. & Yan, Q. Axiomatic Attribution for Deep Networks. in Proceedings of the 34th International Conference on Machine Learning (eds. Precup, D. & Teh, Y. W.) vol. 70 3319–3328 (PMLR, 06--11 Aug 2017).

25. DOI:10.5281/zenodo.8274879.

## References

1. Mostafavi, S., et al. A molecular network of the aging human brain provides insights into the pathology and cognitive decline of Alzheimer’s disease. Nat. Neurosci. 21, 811–819 (2018).

2. Loh, P.-R., et al. Reference-based phasing using the Haplotype Reference Consortium panel. Nat. Genet. 48, 1443–1448 (2016).

3. Novakovsky, G., Dexter, N., Libbrecht, M. W., Wasserman, W. W. & Mostafavi, S. Obtaining genetics insights from deep learning via explainable artificial intelligence. Nat. Rev. Genet. (2022) doi:10.1038/s41576-022-00532-2.

4. Zhou, J. & Troyanskaya, O. G. Predicting effects of noncoding variants with deep learning-based sequence model. Nat. Methods 12, 931–934 (2015).

5. Sundararajan, M., Taly, A. & Yan, Q. Axiomatic Attribution for Deep Networks. in Proceedings of the 34th International Conference on Machine Learning (eds. Precup, D. & Teh, Y. W.) vol. 70 3319–3328 (PMLR, 06--11 Aug 2017).

6. Simonyan, K., Vedaldi, A. & Zisserman, A. Visualising image classification models and saliency maps. Deep Inside Convolutional Networks.

